# Tumor-derived exosomal miR-222-3p induce cancer-associated fibroblasts activation to foster progression of renal cancer

**DOI:** 10.1101/2023.06.22.546057

**Authors:** Yang Yang, Jie Zhu, Dong-lai Shen, Ce Han, Chen-feng Wang, Bo Cui, Wen-mei Fan, Yan Huang, Xiu-bin Li, Xu Zhang, Yu Gao

## Abstract

The interaction between tumor-derived exosomes and stroma is crucial for tumor progression. However, the mechanisms by which tumor cells influence stromal changes are not yet fully understood. Our study revealed that high-metastatic renal cancer cells are more effective in converting normal fibroblasts into cancer-associated fibroblasts (CAFs) compared to low-metastatic renal cancer cells. Meanwhile, high-metastatic renal cancer cells secrete more exosomal miR-222-3p, which can directly target PANK3, activate NF-kB signaling pathway in fibroblasts and induce intracellular metabolic reprogramming to produce more lactic acid through Warburg effect. The activated CAFs further promote renal cancer progression by secreting lactic acid and inflammatory cytokines, including IL-6 and IL-8. Patients with renal cancer who have high levels of serum exocrine miR-222-3p are more likely to experience progression. These findings suggest that the intercellular communication between renal cancer cells and fibroblasts is facilitated by tumor exosomes. Targeting this communication may hold promise for the prevention and treatment of renal cancer.

## 1. Introduction

Pulmonary metastasis and tumor thrombosis are the most common aggressive progression of clear cell renal cell carcinoma (ccRCC) and one of the leading causes of cancer-related death^[1-2]^, and this happens due to the communication between tumor and stromal cells in the microenvironment^[3-4]^. In the fight against tumor progression, therapeutic strategies targeting microenvironment components have gained importance in recent years^[5-9]^. Among the various cell types in the tumor stroma, cancer-associated fibroblasts (CAFs) are the most abundant and play a crucial role in promoting tumor progression and metastasis^[10]^. Due to the fact that CAFs are not a uniform cell type, they exhibit a high level of heterogeneity and can express various specific markers for identification. Two commonly used markers of CAFs are α-smooth muscle actin (α-SMA) and fibroblast activation protein (FAP)^[11]^. Additionally, CAFs have the ability to regulate the inflammatory microenvironment by expressing proinflammatory genes^[12]^. The interaction between tumor cells and CAFs has been extensively researched in ccRCC. However, the specific mechanism through which tumor cells activate fibroblasts is not yet clearly understood, even less in ccRCC progression.

First discovered in the 1980s, exosomes are believed to be bilayer vesicles with a diameter ranging from 50 to 150 nm^[13-14]^. These tiny structures can be produced by various cell types and are released into the extracellular environment through fusion with the cell membrane. Exosomes are identified by the presence of highly enriched specific proteins like TSG101, CD63, HSP70, CD9, and CD81^[15-17]^. These vesicles have been found to play crucial roles in the transmission of genetic material, such as nucleic acids, activation of signaling pathways, and metabolic reprogramming. They also have significant implications in tumor progression, response to drugs and immune cells^[18-20]^.

MicroRNAs (miRNAs) are small, non-coding RNAs that inhibit the translation of target mRNAs^[21]^. Recent research has revealed that exosomes contain high levels of miRNAs, which are crucial in regulating the immune system, drug resistance, and metastasis of various tumor types^[22-24]^. However, the relationship between tumorigenic exosomal miRNAs and the progression of ccRCC remains unclear. This study aims to investigate the causes of ccRCC progression through intercellular communication. High and low metastatic renal cancer cell lines were compared using exosomes microarray comparison, leading to the identification of tumor-derived exogenous miRNAs that convert fibroblasts into CAFs. Furthermore, CAFs were found to promote tumor progression by increasing the secretion of lactic acid and inflammatory cytokines. The crosstalk between stromal and tumor cells revealed a new mechanism of ccRCC progression and provides a potential therapeutic strategy for advanced ccRCC, including lung metastasis and tumor thrombosis.

## 2. Materials and methods

### Specimens and primary cells

A tissue microarray comprising of localized or progressive ccRCC tissue samples from 206 patients was obtained from the Chinese PLA General Hospital. The clinicopathological features of the patients are shown in Supplementary Table 1. Primary isolated CAFs were cultured in DMEM/F12 supplemented with 10% FBS (Pricella). All procedures were approved by the Ethics Committee of Chinese PLA General Hospital and informed consent was obtained before the study.

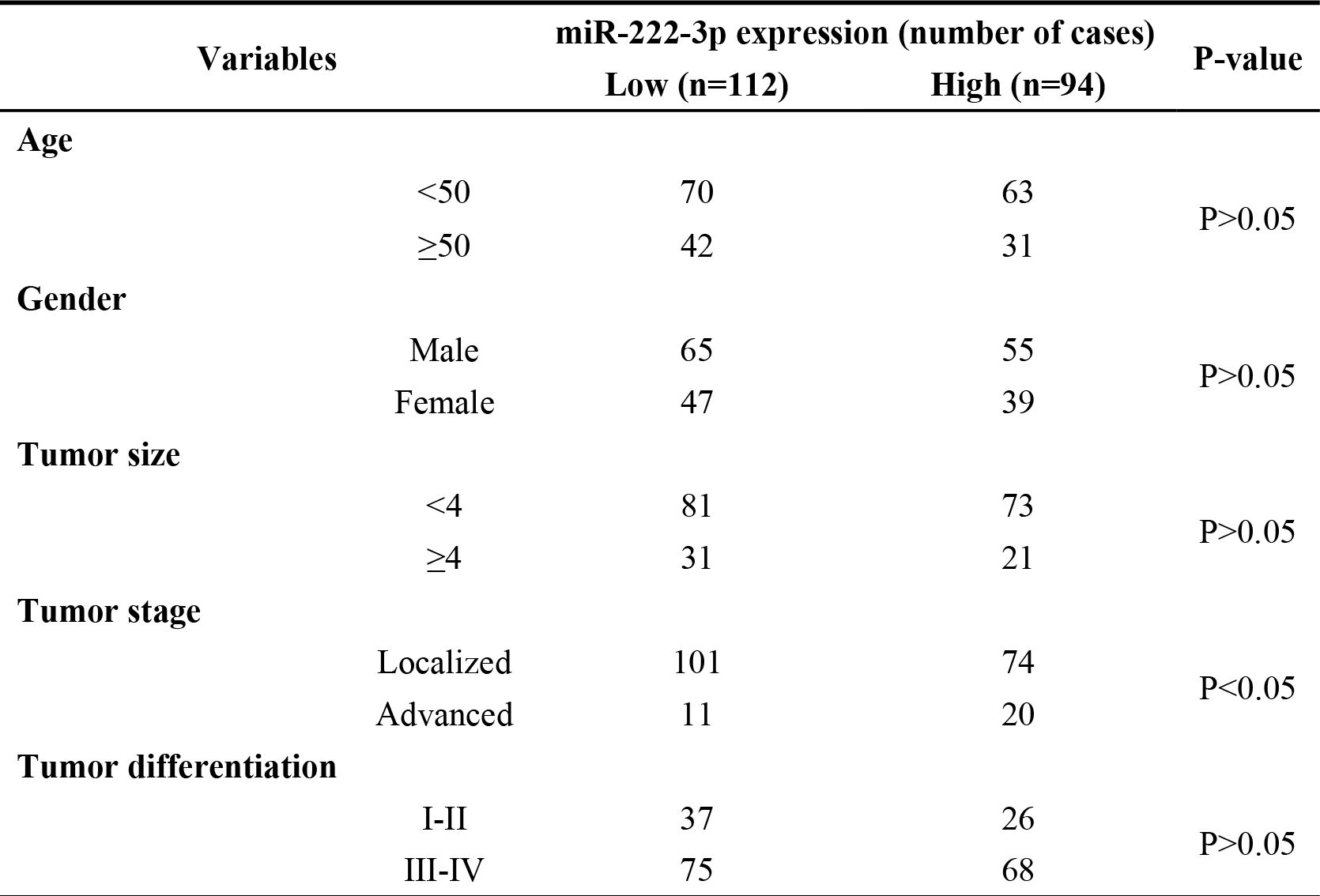
**Supplementary Table 1** Summary of Clinicopathologic Variables.

### Cell culture

Human renal carcinoma cell lines OSRC2, 786-O, ACHN, and SN12 were obtained from the Cell Bank of Type Culture Collection of the Chinese Academy of Sciences. The cells were cultured in 1640, 1640, MEM, and DMEM (Pricella), respectively, with the addition of 10% FBS (Pricella). The human renal fibroblast cell line (NFs) was obtained from Pricella Company (CP-H072) and cultured in a specialized medium (Pricella, CM-H072) at 37℃ in a humidified incubator with 5% CO_2_. The cell lines were identified using short tandem repeats (STR) analysis and confirmed to be free of mycoplasma contamination.

### Reagents and antibodies

Antibodies for TSG101 (ab125011, 1 : 1000), CD81 (ab79559, 1 :1000), α-SMA (ab7817, 1 : 2000) were purchased from Abcam. Antibody for CD63 (A5271, 1 : 1000) was purchased from ABclonal. Antibodies for FAP (66562s, 1 : 1000), Phosphotylated NF-kB (3033, 1:1000), NF-kB (8242, 1:1000), IkBα (4812, 1:1000) and GAPDH (5174S, 1:1000) were purchased from Cell Signaling Technology. Antibody for PANK3 (bs-8339R, 1 : 1000) was purchased from Bioss. Antibodies for MCT1 (abs120479, 1 : 1500) and MCT4 (abs124388, 1 : 2000) were purchased from Absin.

### Western blotting

Whole-cell protein extracts were homogenized in cell lysates and centrifuged at 12000 rpm for 15 minutes to obtain the protein concentration, which was measured using the bicinchoninic acid (BCA) assay. The protein was then transferred onto a nitrocellulose filter membrane and incubated with a specific antibody. The immune complex was subsequently incubated with the corresponding secondary antibody and detected using Tanon chemiluminescence imaging system.

### RNA interference and plasmids

siRNAs (siNC, siRNA targeting miR-222-3p or PANK3) and mimics of indicated miRNAs were conducted by Ribobio Company (Guangzhou, China). The sequences of siRNAs and miRNA mimic referred above were listed in Supplementary Table 2-3. The plasmid vector containing PANK3 and empty vector were provided by Omiget Technology (Beijing, China).

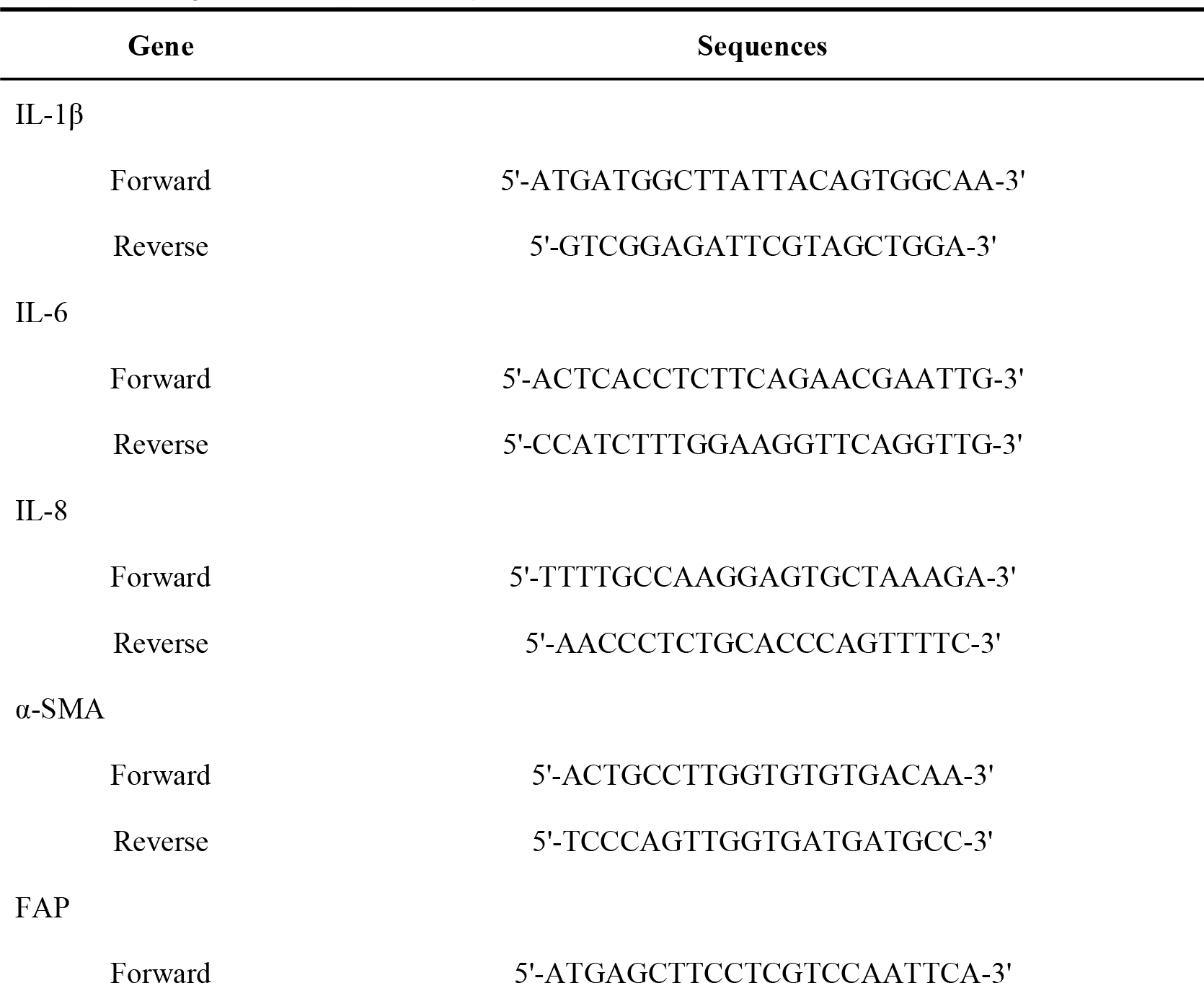
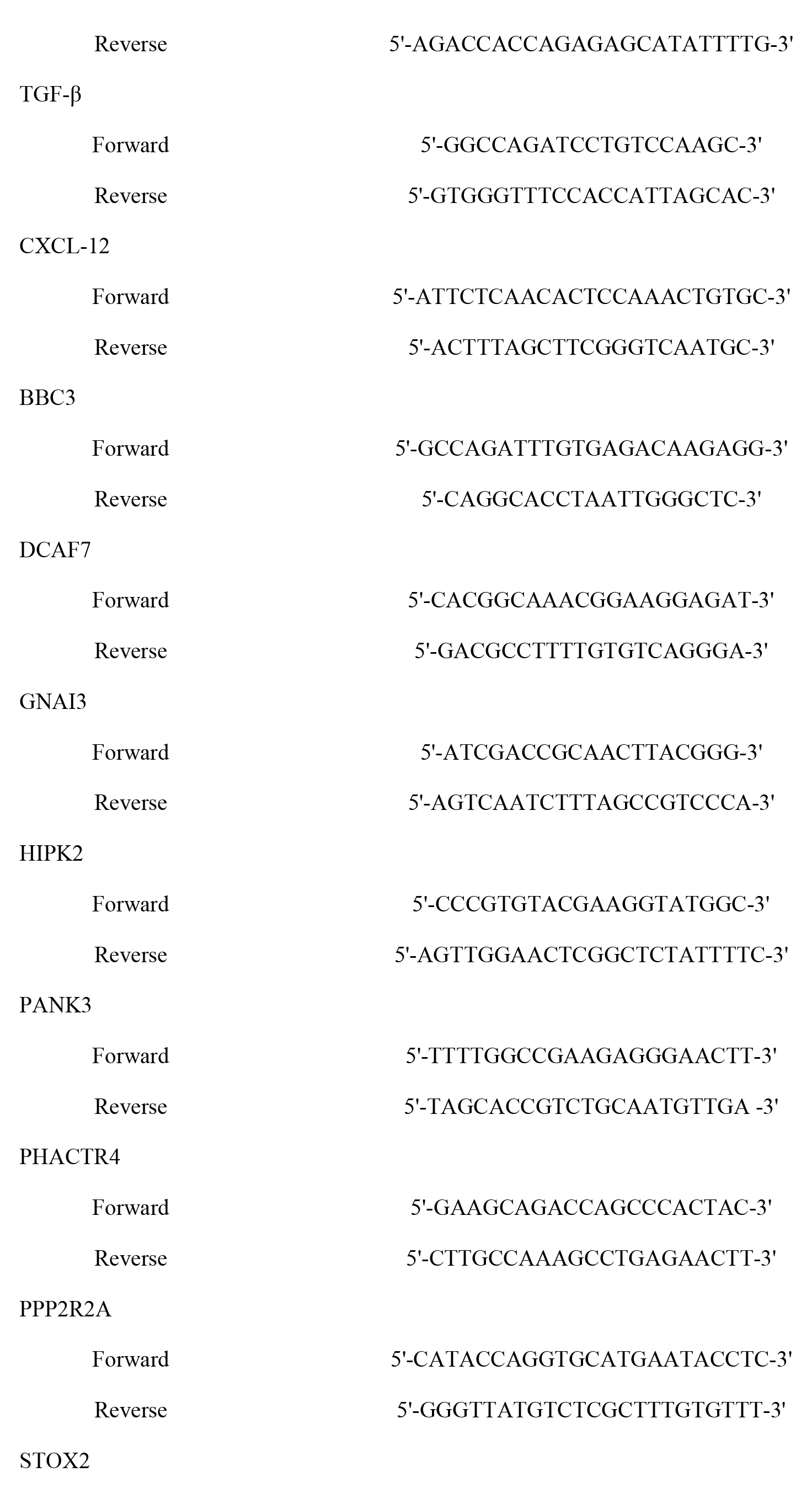
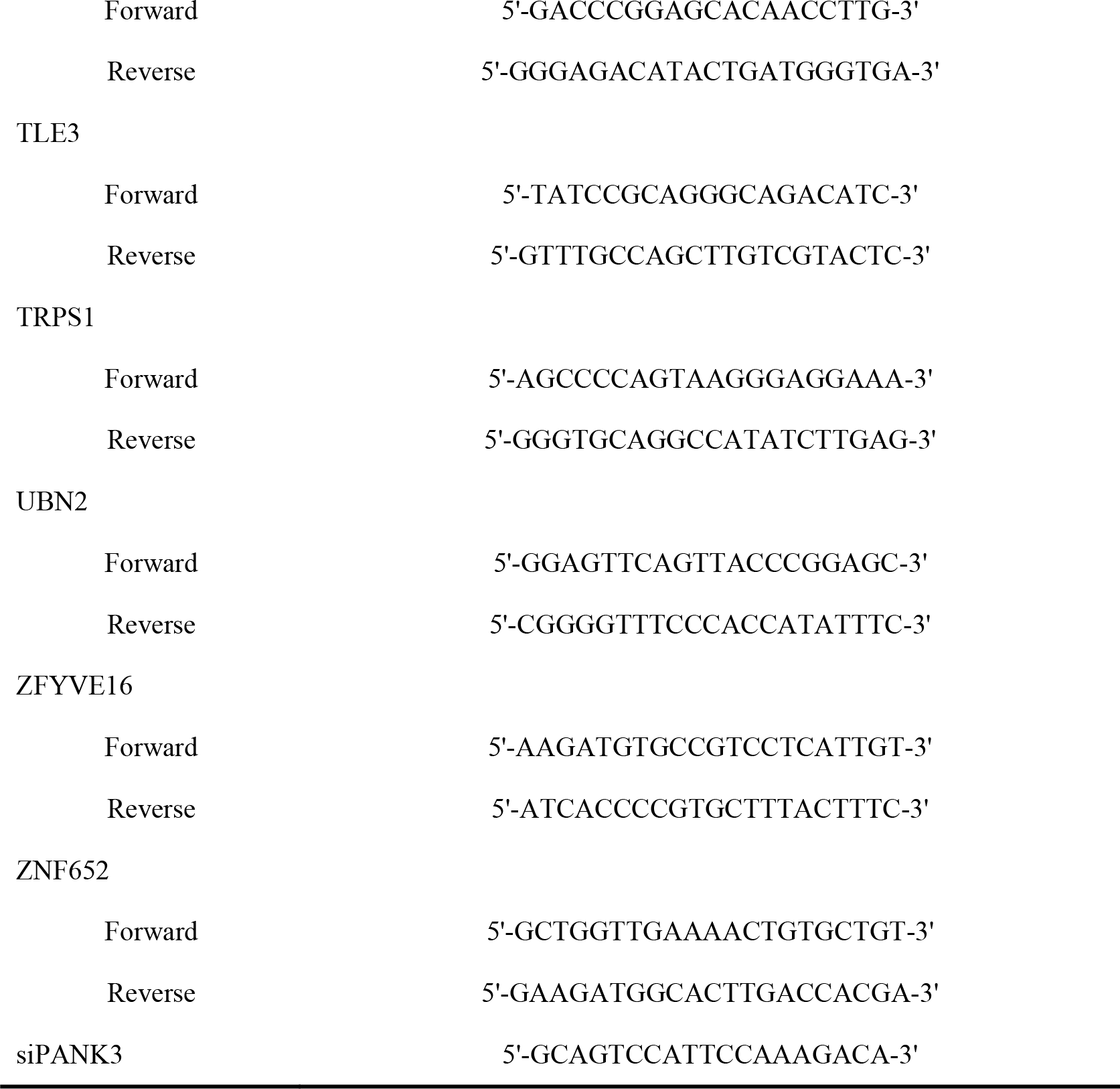
**Supplementary Table 2** Sequences of primers and siRNA.

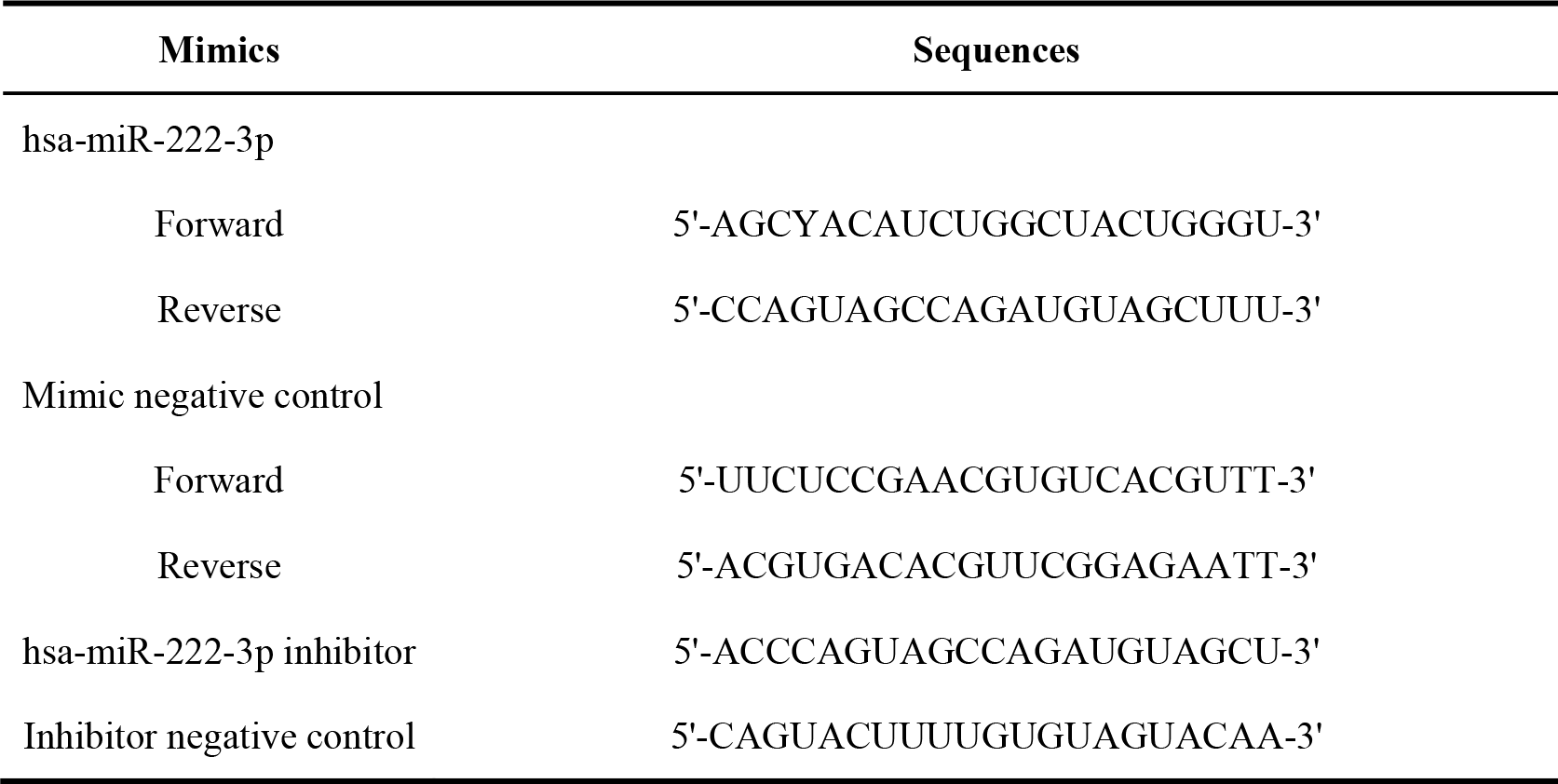
**Supplementary Table 3** Sequences of miRNA mimics.

### Animal studies

In order to study the impact of exosomes on lung metastasis, we injected 1×10^6^ luciferase-labeled SN12 cells into 4-week-old male nude mice via tail vein. The mice were then randomly divided into groups and given intravenous injections of exosomes from indicated carcinoma cell line conditioned mediums twice a week for 6 weeks. The Tanon imaging system was utilized to analyze the resulting lung metastasis.

For xenograft assays, 1×10^6^ educated SN12 cells were injected into the renal capsule of nude mice. The size of the tumors was measured at specific times. All animal experiments were approved by the Committee on Use and Care of Animals of Chinese PLA General Hospital.

### Migration assay and wound-healing assay

For migration assay, Matrigel diluent was used to coat the upper surface of the membrane at the bottom of the Transwell chamber, and the cells prepared were cultured in serum-free medium for 12 h in advance. The cell density was adjusted to 1×10^5^ / ml. Add 500μl complete medium to the lower chamber of Transwell, and add 200μl cell suspension prepared in front of the upper chamber. The cell migration ability was cultured in a CO_2_ incubator. Staining was performed using crystal violet solution, and the number of cells was averaged by randomly selecting three fields.

For wound-healing assay, cells were seeded in 6-well plates at 80% density. A horizontal line was drawn using the tip of a pipette, and the complete medium corresponding to the cells was added along with mitomycin. The plates were then placed in a CO_2_ incubator, and at a fixed time, the same field of view was selected under a bright field microscope to photograph and quantitatively analyze the speed and area of the scratch covered.

### RNA extraction and qRT-PCR

Total RNA was extracted using RNA-Quick Purification Kit (ES science, RN001) according to the manufacturer’s instructions. Reverse transcription was performed using Fast All-in-One RT Kit (ES science, RT001). Real-time PCR was conducted using Super SYBR Green qPCR Master Mix (ES science, QP002). For miRNAs, reverse transcription was performed using microRNA Reverse Transcription Kit (ES science, EZB-miRT4-L). Real-time PCR was conducted using EZ-Probe qPCR Master Mix for microRNA (ES science, EZB-miprobe-R2). The sequences of all indicated primers were listed in Supplementary Table 2.

### Immunohistochemistry and in situ hybridization analysis

For immunohistochemistry, the slides were initially incubated with primary antibodies. Subsequently, the slides were subjected to procedures utilizing the Immunohistochemistry Kit (Biosharp) in accordance with the manufacturer’s instructions. For in situ hybridization analysis, the slides underwent treatment using the hsa-miR-222-3p miRCURY LNA detection probe and the Enhanced Sensitive IDH Detection Kit (Boster). The resulting images were captured and assessed using TissueFAXS software.

### Isolation and analysis of exosomes

Exosome-free serum was incorporated into the cell culture medium configuration. Exosomes were extracted from the conditioned medium through traditional ultra-speed differential centrifugation. The exosomes were then observed through transmission electron microscopy and quantified using Nano-Sight NS300.

### Luciferase reporter assay

Ribobio Company designed and synthesized the dual luciferase reporter vector with the renilla luciferase gene (hRluc) as the reporter fluorescence and the firefly luciferase gene (hluc) as the corrected fluorescence. The 3’UTR region of the gene was cloned downstream of the hRluc gene, and the miRNA was co-transfected with the reporter gene vector. The down-regulation of the relative fluorescence value of the reporter gene confirmed the interaction between the miRNA and the target gene.

### Measurement of lactate production, OCR and ECAR

Lactate Colorimetric Assay Kit (Sigma-Aldrich) was used according to the manufacturer’s instructions.The Seahorse Bioscience XFe96 Extra-cellular Flux Analyzer platform was employed to analyze the mitochondrial oxygen consumption rate (OCR) and extracellular acidification rate (ECAR). The Seahorse software was utilized to analyze the results.

## 3. Results

### Tumor-derived exosomes regulate fibroblasts activation

In the metastatic niche, research has shown that cancer-associated fibroblasts (CAFs) play an active role in promoting tumor metastasis progression^[25]^. Within the CAFs subgroup, those expressing α-SMA are considered particularly aggressive^[11]^. As a result, it is important to investigate the communication between cells, specifically how tumor cells activate fibroblasts. To investigate whether exosomes play a role in activating fibroblasts, given their importance in inducing systemic changes in tumor tissue. Four renal cancer cell lines were selected, including high-metastatic cell lines SN12 and ACHN, and low-metastatic cell lines OSRC2 and 786-O. Exosomes were extracted from the conditioned medium using ultracentrifugation and were observed to have a ’teacup shape’ under transmission electron microscope (Fig. 1A). We utilized Nanosight particle tracking analysis to determine that the exosomes from indicated conditioned medium were mostly within 100-150nm range. The concentration of exosomes with a high-metastatic background were found to be enriched at a higher level than those with a low-metastatic background(Fig.1B). Additionally, the presence of characteristic exosomal markers CD63, CD81, and TSG101 provided further confirmation that the isolated particles were indeed exosomes(Fig.1C).

**Fig. 1.**
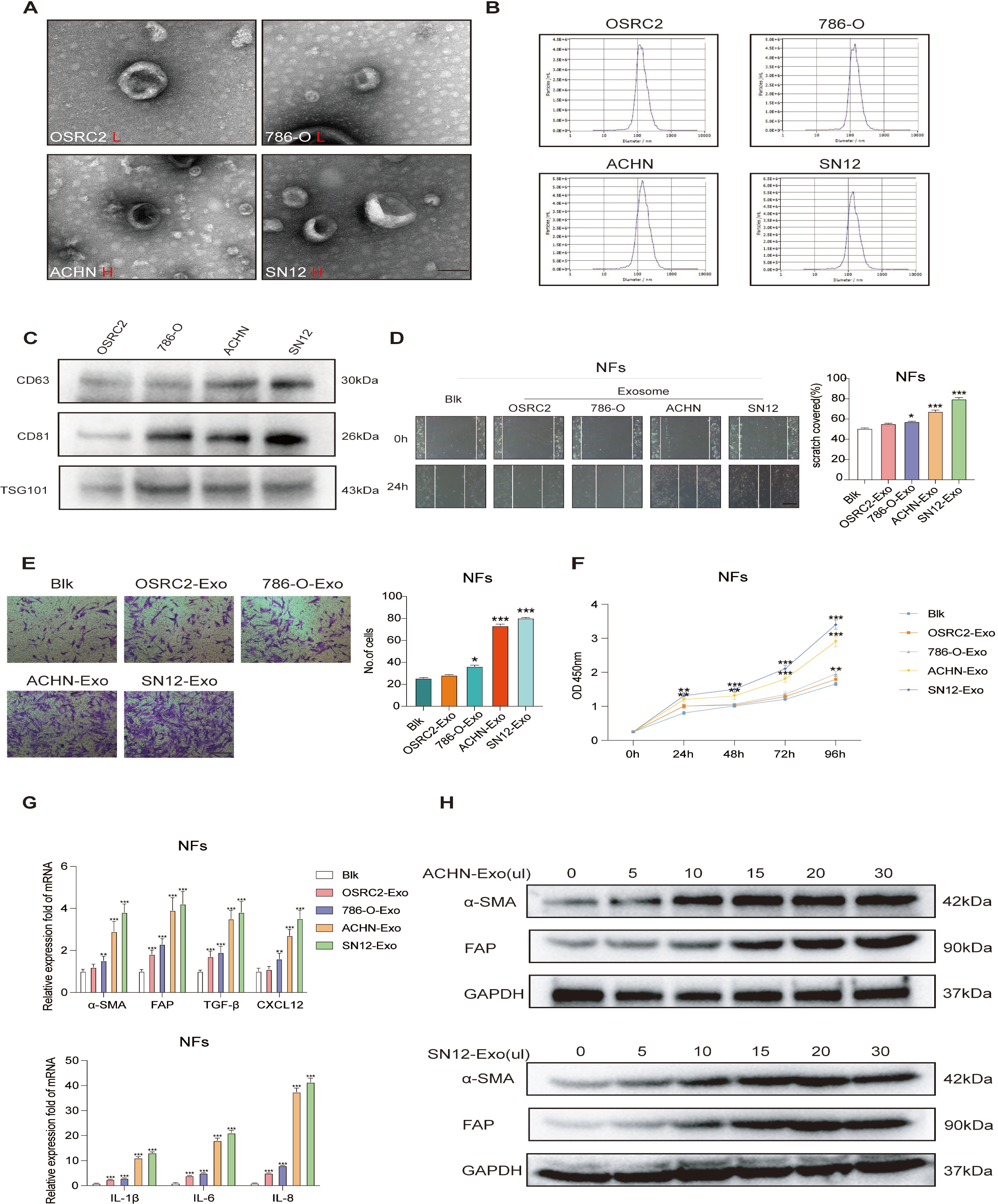
Exosomes secreted from high-metastatic renal cancer cells regulate fibroblasts activation. **A-B** Electron microscopy and Nanosight particle tracking analysis of exosomes released by different renal cancer cell lines. Scale bar, 100 nm. **C** Western blotting assay of indicated proteins in exosomes from different renal cancer cells. **D-F** Wound-healing,migration and CCK-8 assays of NFs treated with equal quantities of exosomes derived from indicated renal cancer cells or blank control. Scale bar, 100 μm. **G** qRT-PCR analysis of indicated gene expression of NFs treated with exosomes released by different renal cancer cells or blank control. **H** Western blotting assay of α-SMA and FAP in NFs co-culture with different concentrations of exosomes from ACHN and SN12 cell lines. Each experiment was performed three times independently and results are presented as mean ± s.d. Student’s t-test was used to analyze the data. (*p < 0.05; **p < 0.01; ***p < 0.001)

Human renal fibroblasts were chosen as normal fibroblasts (NFs) and subjected to further evaluation of the effects of exosomes derived from four types of conditioned medium. Wound-healing and transwell assays were performed, and the results indicated that exosomes from high-metastatic cell lines significantly improved the migration and proliferation abilities of the fibroblasts(Fig1D-F). Additionally, co-culturing NFs with exosomes for 48 hours led to the upregulation of CAFs markers (α-SMA and FAP) and proinflammatory genes(IL-1β, IL-6, IL-8, TGF-β, and CXCX12)(Fig1G). These findings suggest that exosomes from high-metastatic cell lines have a significant impact on the activation of fibroblasts. CXCL12, also known as stromal cell derived factor −1 (SDF-1), enhances the mobility of stromal fibroblasts, particularly myofibroblasts^[26]^. The aforementioned findings demonstrate that exosomes can induce fibroblasts to express pro-inflammatory genes at high levels, leading to the formation of an inflammatory microenvironment and promoting the activation of fibroblasts into CAFs. Furthermore, the concentration of exosomes plays a crucial role in the activation of fibroblasts. As the concentration of exosomes increased, the expression levels of α-SMA and FAP proteins in fibroblasts also increased. However, after adding more than 15μl of exosome solution, the expression levels remained stable(Fig1H), suggesting that high-metastatic background exosomes have the ability to activate fibroblasts.

### Exosomal miR-222-3p mediates fibroblasts activation

We next investigated the activation of fibroblasts by tumor-derived exosomes. Proposing that miRNAs, which are small endogenous non-coding RNAs expressed in cells^[22-23]^, were responsible for mediating fibroblast activation. To identify the target miRNA, we conducted microarray analysis on exosomal miRNAs from four renal cancer cell lines. The results revealed that 14 miRNAs were upregulated (fold change > 10) in both high-metastatic renal cancer cell-derived exosomes. These miRNAs were subjected to validation and miR-222-3p was found to be the most significant(Fig2A-B). This result was consistent with the differential expression of miRNAs in blood exosomes of 4 patients with localized ccRCC, 5 ccRCC patients with inferior vena cava tumor thrombus and 4 ccRCC patients with lung metastasis in our department(Fig2C). Quantitative reverse-transcriptase PCR (qRT-PCR) analysis further confirmed the level of miR-222-3p in exosomes from different metastatic background renal tumor cell lines. The findings revealed that the level of miR-222-3p in exosomes from cell lines with metastatic background was significantly higher compared to those without. These results suggest that the level of miR-222-3p may be linked to the progression of ccRCC (Fig2D). To explore the correlation between miR-222-3p levels and clinicopathological features of ccRCC, we conducted miR-222-3p in situ hybridization on 206 localized or progressive ccRCC tissues. The specimens were then categorized into two groups based on their miR-222-3p expression score, namely low and high miR-222-3p expression groups. According to follow-up results, high miR-222-3p expression predicted poor survival outcomes, which was consistent with TCGA database results(Fig2E). To further validate the results, we transfected miR-222-3p mimic and inhibitor into NFs and verified the levels of miR-222-3p through qRT-PCR (Fig2F). Importantly, miR-222-3p mimic had a significant impact on the expression of proinflammatory genes, including IL-1β, IL-6, IL-8, TGF-β, and CXCL12 in NFs. Additionally, the mimic was found to enhance the expression of CAFs markers α-SMA and FAP in fibroblasts(Fig2G). Wound-healing and transwell assays demonstrated that the transfection of miR-222-3p mimic significantly improved the migration ability of fibroblasts(Fig2H). Further more, the CCK-8 assay indicated that miR-222-3p mimic enhanced the proliferation of fibroblasts(Fig2J).

**Fig. 2.**
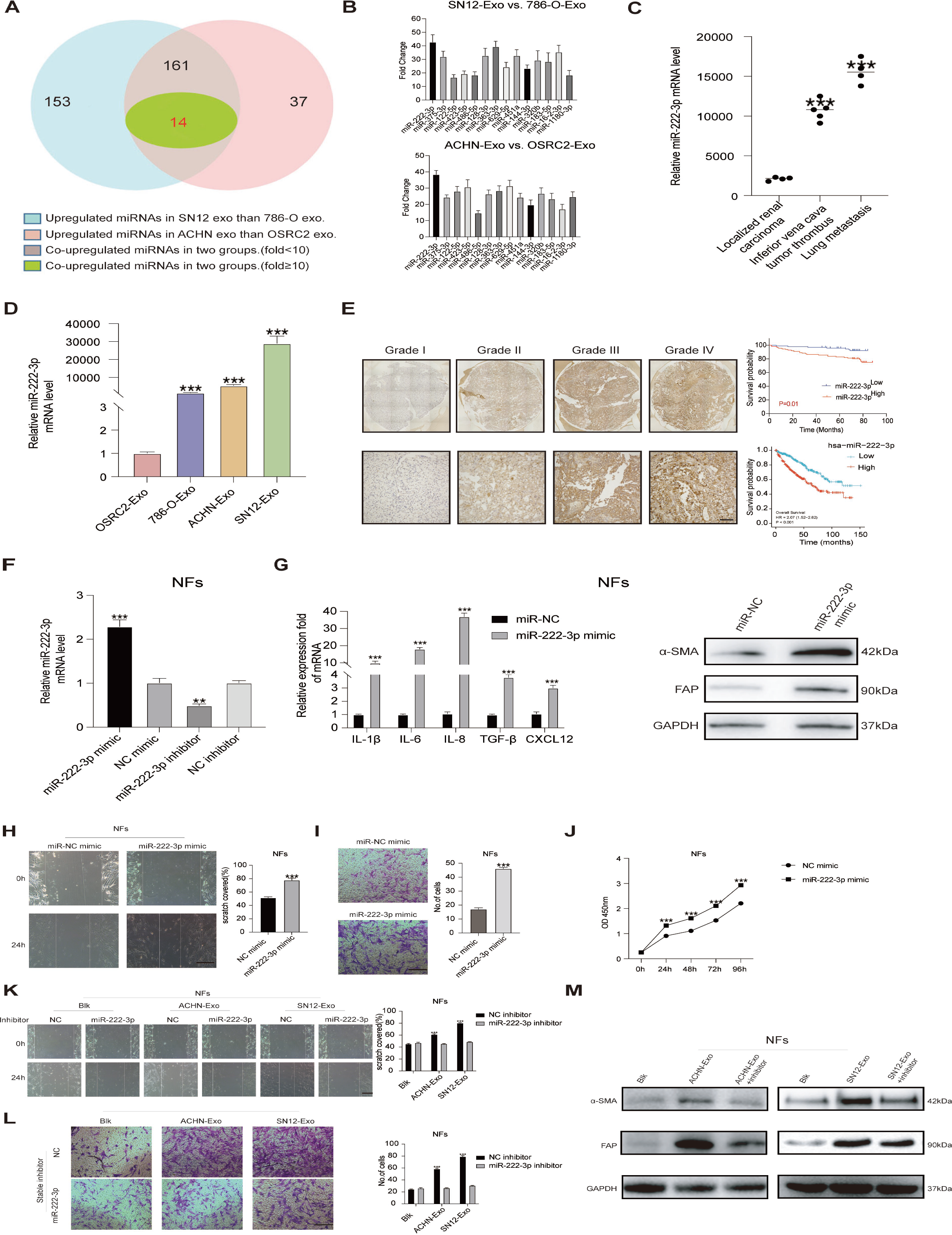
Exosomal miR-222-3p is characteristically secreted by high-metastatic renal cancer cells and mediates fibroblasts activation. **A-B** Overlapping results of upregulated miRNAs in indicated groups. **C** Blood-derived exosomal miR-222-3p expression in indicated patient groups. **D** Exosomal miR-222-3p expression from different renal cancer cells. **E** In situ hybridization of miR-222-3p on serial sections of human RCC tissues. Scale bar, 50 μm. **F** qRT-PCR assay of miR-222-3p expression in NFs transfected with miR-222-3p mimic or inhibitor. **G** qRT-PCR and western blotting assays of indicated proinflammatory gene and CAFs markers (α-SMA and FAP) expression of NFs transfacted with miR-222-3p mimic or normal control. **H-J** Wound-healing,migration and CCK-8 assays of NFs transfacted with miR-222-3p mimic or normal control. Scale bar, 100 μm. **K-L** Migration ability comparison of NFs treated with exosomes derived from ACHN or SN12 with stably expressing miR-222-3p inhibitor or negative control. Scale bar, 100 μm. **M** Western blotting assay of α-SMA and FAP expression of NFs treated with exosomes derived from ACHN or SN12 with stably expressing miR-222-3p inhibitor or negative control. Each experiment was performed three times independently and results are presented as mean ± s.d. Student’s t-test was used to analyze the data. (*p < 0.05; **p < 0.01; ***p < 0.001)

To further investigate the effect of miR-222-3p, fibroblasts cocultured exosomes were transfected with miR-222-3p inhibitors. The expected outcome was observed as the effect of exosomes on fibroblasts was attenuated by its specific inhibitors(Fig2K-M). These findings collectively reveal that tumor-derived exosomal miR-222-3p mediate fibroblast activation.

### Exosomal miR-222-3p directly targets PANK3 in fibroblasts

In order to investigate the target gene of miR-222-3p, we utilized four bioinformatics tools (miRDB, miRWalk, TargetScan, and miRTarBase) to predict common target genes. Our analysis revealed 13 target gene intersections(Fig3A). The relative changes of these 13 target genes of NFs transfected with miR-222-3p mimic were confirmed through qRT-PCR(Fig3B). Combined with the differential expression and survival prognosis analysis of TCGA database, we finally selected pantothenate kinase 3 (PANK3) as a possible downstream target gene(Fig3C). First, we confirmed that miR-222-3p reduces the expression of PANK3expression in NFs(Fig3D). Then we compared the PANK3 full-length sequence with the miR-222-3p sequence and found that the PANK3 coding sequence could be the target of miR-222-3p. To confirm this, the wild type and mutant miR-222-3p binding sites were cloned into luciferase vectors. The results showed that in the presence of miR-222-3p, the luciferase activity decreased significantly in cells transfected with the wild binding site vector NFs. However, cells with a mutant binding site vector did not show such repression(Fig3E). The results indicate that miR-222-3p directly targets PANK3. siPANK3 and PANK3 overexpression plasmids were separately transfected into NFs, and the levels of PANK3 were verified through qRT-PCR and western blot analysis(Fig3F-G). The migration ability of NFs was observed to increase upon transfection with siRNAs targeting PANK3(Fig3H). In addition, overexpression of full-length PANK3 can counteract the impact of miR-222-3p on fibroblast migration(Fig3I). Moreover, when fibroblasts were transfected with siPANK3, the expression of pro-inflammatory genes such as IL-1β, IL-6, and IL-8 increased(Fig3J), and this effect was reversed when PANK3 overexpression plasmids were subsequently transfected(Fig3K).

Immunofluorescence results indicated that PANK3 expression was significantly lower in ccRCC tissues compared to adjacent normal renal tissues, while FAP levels were significantly increased, suggesting that PANK3 levels in CAFs may be low(Fig3L). To further confirm PANK3 expression in CAFs, we isolated two CAFs from tissue samples of patients with advanced ccRCC. Results showed a significant decrease in PANK3 protein levels in CAFs compared to NFs(Fig3M). Overall, these findings suggest that miR-222-3p directly targets PANK3 and mediates fibroblast activation.

**Fig. 3.**
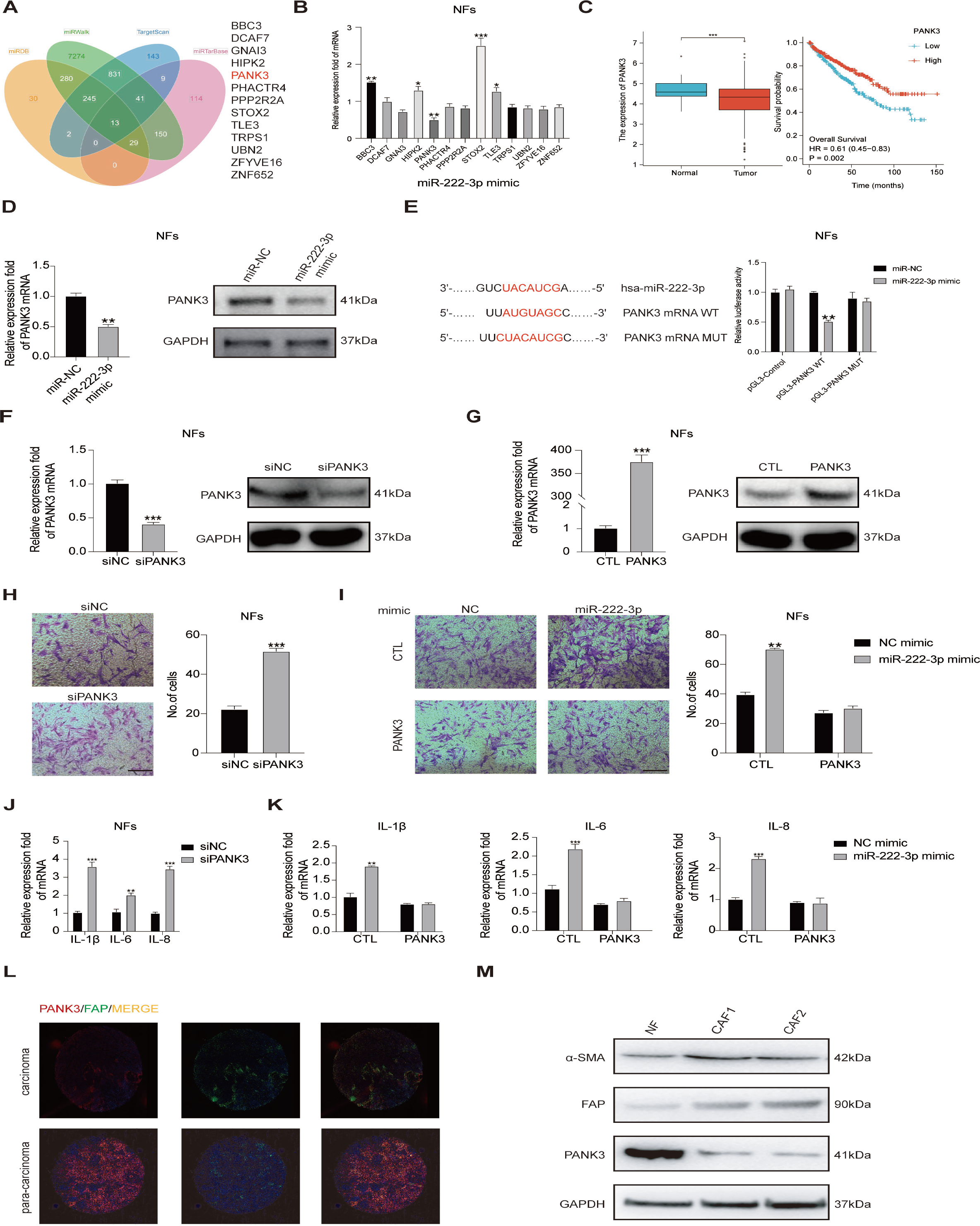
Exosomal miR-222-3p directly targets PANK3 in fibroblasts. **A** Target gene prediction of miR-222-3p with four bioinformatics tools. **B** qRT-PCR assay of relative fold changes of target genes expression in NFs transfected with miR-222-3p mimic. **C** Differential expression and prognostic analysis of PANK3 in TCGA database. **D** qRT-PCR and immunoblotting assays of PANK3 expression in NFs transfected with miR-222-3p mimic or normal control. **E** Relative luciferase activity of NFs in the presence of indicated treatments. **F** qRT-PCR and immunoblotting assays of PANK3 expression in NFs transfected with siRNAs targeting PANK3 or control. **G** qRT-PCR and immunoblotting assays of PANK3 expression in the presence of PANK3 or not. **H** Migration assay of NFs transfected with siRNAs targeting PANK3 or control. Scale bar, 100 μm. **I** MiR-222-3p effect on migration ability of NFs in the presence of PANK3 or not. Scale bar, 100 μm. **J** qRT-PCR analysis of pro-inflammatory genes expression in NFs transfected with siRNAs targeting PANK3 or control. **K** qRT-PCR analysis of pro-inflammatory genes expression in NFs with indicated treatments. **L** Immunofluorescence detection of PANK3 and FAP in renal cell carcinoma and para-carcinoma tissues. **M** Western blotting assay of indicated proteins in NFs and CAFs. Each experiment was performed three times independently and results are presented as mean ± s.d. Student’s t-test was used to analyze the data. (*p < 0.05; **p < 0.01; ***p < 0.001)

### MiR-222-3p activation of fibroblasts via PANK3-NF-kB axis and metabolic reprogramming

Transcriptome sequencing of NFs transfected with si PANK3 and si NC revealed significant changes in the expression levels of multiple genes involved in the NF-kB signaling pathway(Fig4A). Our results indicate that miR-222-3p enhances the expression of proinflammatory genes in NFs, including well-known targets of NF-kB such as IL-1β, IL-6, and IL-8. Furthermore, exosomes from high-metastatic tumor cells were found to promote the expression of p-NF-kB in NFs, inhibit IkBα, and activate the NF-kB signaling pathway(Fig4B). These findings suggest a potential role of miR-222-3p and exosomes in regulating NF-kB signaling in tumor metastasis. In addition, the transfection of miR-222-3p mimic or si PANK3 led to the promotion of NF-kB phosphorylation and inhibition of IkBα, whereas overexpression of PANK3 inhibited NF-kB phosphorylation(Fig4C). These findings suggest that miR-222-3p activates fibroblasts by mediating the activation of NF-kB signaling.

**Fig. 4.**
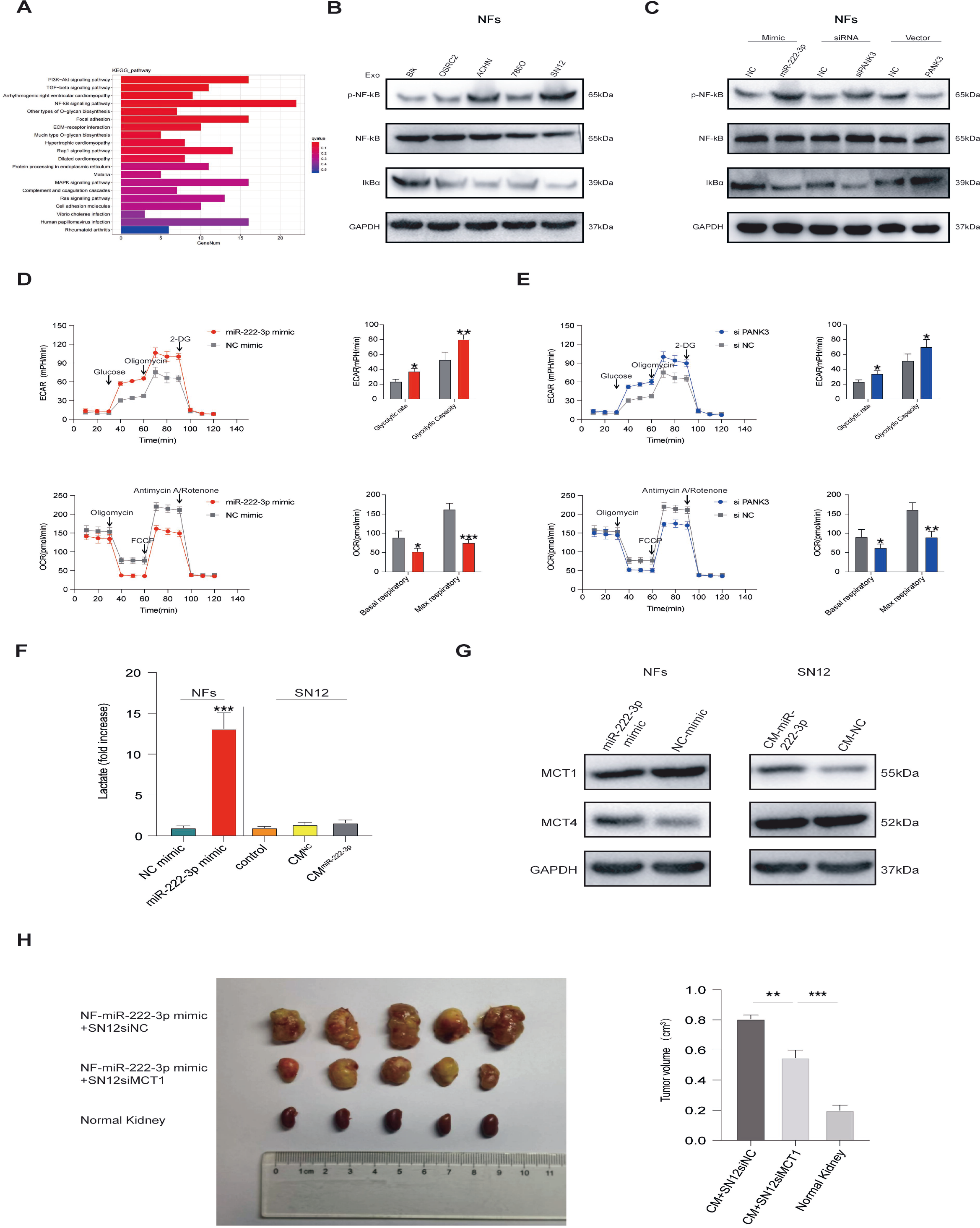
MiR-222-3p activation of fibroblasts via PANK3-NF-kB axis and metabolic reprogramming. **A** KEGG enrichment analysis of differentially expressed genes between NFs transfected with siRNAs targeting PANK3 or control. **B-C** Immunoblotting assays of indicated proteins in NFs with indicated treatments. **D-E** Analysis of ECAR and OCAR in NFs with indicated treatments. **F** Detection of lactic acid in the conditioned medium of NFs or SN12 with indicated treatments. **G** Immunoblotting assays of MCTs in NFs and SN12 with indicated treatments. **H** Xenograft assays of SN12 with indicated treatments were performed on nude mice. Representative tumors (left) and tumor volumes (right) were shown. Each experiment was performed three times independently and results are presented as mean ± s.d. Student’s t-test was used to analyze the data. (*p < 0.05; **p < 0.01; ***p < 0.001)

PANK3 is an essential enzyme in the de novo synthesis of CoA, controlling the rate of CoA synthesis^[27]^. Our study measured the ECAR and OCR levels of NFs to determine the impact of miR-222-3p and PANK3 on NFs metabolism. The results indicated that the glycolytic rate and capacity of NFs transfected with miR-222-3p mimic and si PANK3 were increased, while the basal and max respiratory levels were decreased. This suggests that the Warburg effect occurs during the activation of fibroblasts(Fig4D-E). Further investigation of the metabolic changes in the activation process of NFs revealed an increase of more than 10 times in the level of lactic acid produced by NFs transfected with miR-222-3p mimic. Then, SN12 cells were cultured in two different types of conditioned medium: one from NFs transfected with miR-222-3p mimic and the other from DMEM serum-free medium. The level of lactic acid in the conditioned medium of NFs transfected with miR-222-3p mimic decreased significantly, suggesting that SN12 cells may consume lactic acid(Fig4F). Monocarboxylate transporters (MCTs) are key molecules in glycolysis, responsible for the transport of lactic acid. MCT4 is mainly responsible for the excretion of intracellular lactic acid, while MCT1 is responsible for the uptake of lactic acid into the cell^[28]^. NFs transfected with miR-222-3p mimic expressed higher levels of MCT4, and SN12 cells co-cultured with conditioned medium of NFs transfected with miR-222-3p mimic expressed higher levels of MCT1. The findings indicate that the activation of fibroblasts leads to a significant production of lactic acid. Additionally, co-cultured SN12 cells are capable of taking up lactic acid into the cell through the MCT1 pathway(Fig4G). In vivo experiments further demonstrated that silencing MCT1 resulted in the inhibition of tumor growth in an orthotopic transplantation tumor model, highlighting the impact of lactic acid uptake on the tumorigenicity of renal cancer cells(Fig4H).

### Activated fibroblasts promote renal cancer progression

To investigate the structural changes of fibroblasts, we utilized transmission electron microscopy to observe NFs, NFs transfected with miR-222-3p mimic, and primary isolated CAF1 and CAF2. The overall and mitochondrial structure of NFs transfected with miR-222-3p mimic, CAF1, and CAF2 were found to be similar. Compared to NFs, the cytoplasm of the aforementioned cells exhibited slight edema, the cell matrix was sparse, the mitochondria were swollen, and the cristae structure was less pronounced(Fig. 5A). Immunohistochemical staining of CAFs marker FAP was performed on three different degrees of ccRCC tissues, including low-grade carcinoma, high-grade carcinoma, and ccRCC with inferior vena cava tumor thrombus in tissue microarray. Our findings revealed that the ratio of CAFs increased in ccRCC tissues with progression, as shown in Fig5B. Furthermore, CAFs-derived conditioned medium was found to promote the migration of renal cancer cells(Fig5C). Notably, CAFs-derived conditioned medium was also found to promote EMT in renal cancer cells and tumorigenesis in vivo(Fig5D-E). In comparison to the NFs group and the control group, the exosomes from CAFs were found to enhance the development of lung metastases caused by SN12 cells. Additionally, it was observed that CAFs were significantly more abundant in metastatic regions. These findings provide further evidence that CAFs play a role in promoting the progression of ccRCC(Fig5F-G).

**Fig. 5.**
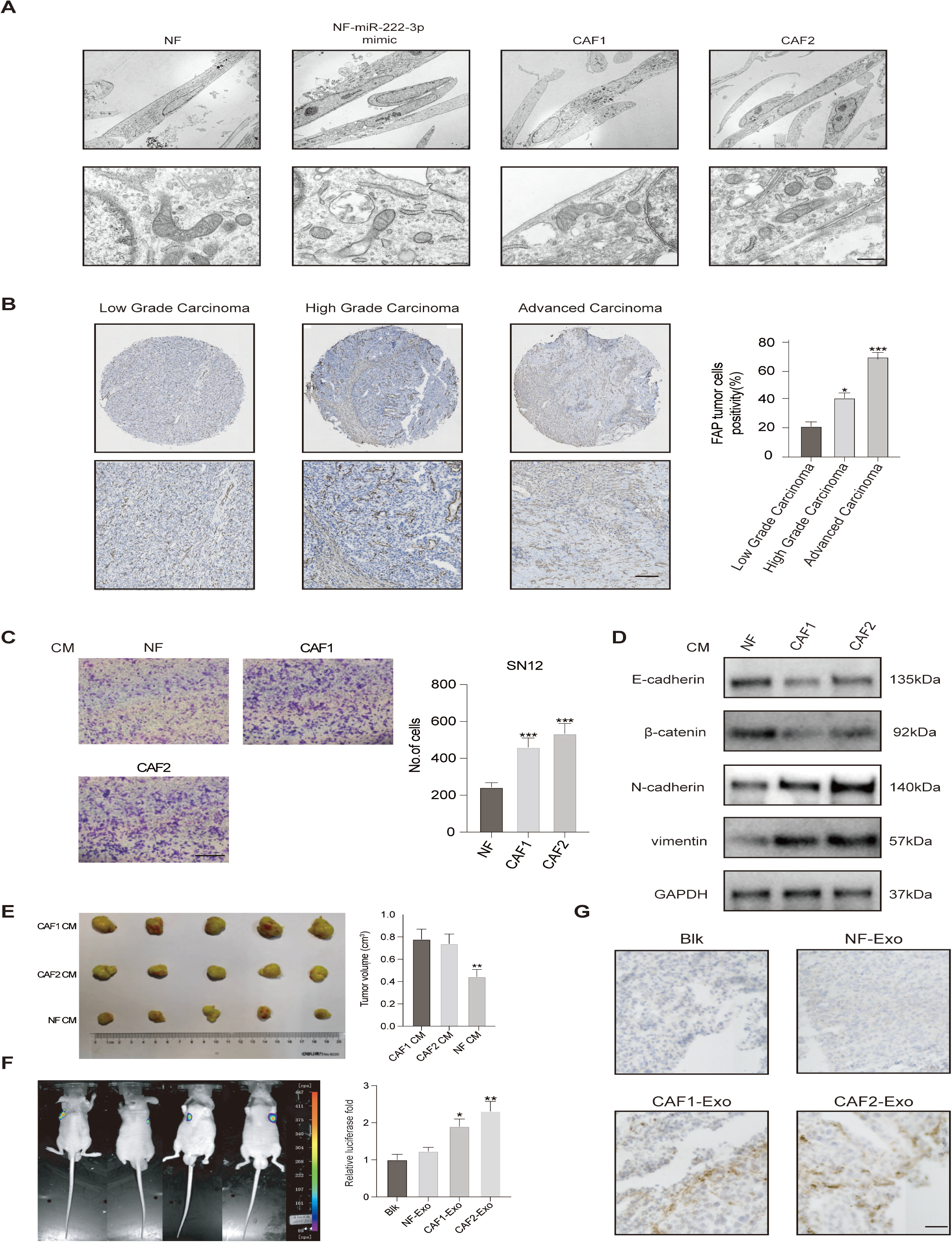
Activated fibroblasts promote renal cancer progression. **A** Transmission electron microscopy images of NFs and CAFs. Scale bar, 50 μm. **B** IHC staining of FAP on serial sections of human RCC tissues. Scale bar, 50 μm. **C** Migration assay of SN12 treated with indicated CM. Scale bar, 100 μm. **D** Immunoblotting assays of EMT markers in SN12 treated with indicated CM. **E** Xenograft assays of SN12 with indicated treatments were performed on nude mice. Representative tumors (left) and tumor volumes (right) were shown. **F** Representative images and quantitative analysis of lung metastasis of indicated mice treated with exosomes derived from NFs or CAFs were determined by luciferase-based bioluminescence imaging. **G** IHC staining of FAP on lung metastatic tissue samples of indicated nude mice. Scale bar, 50 μm.

**Fig. 6.**
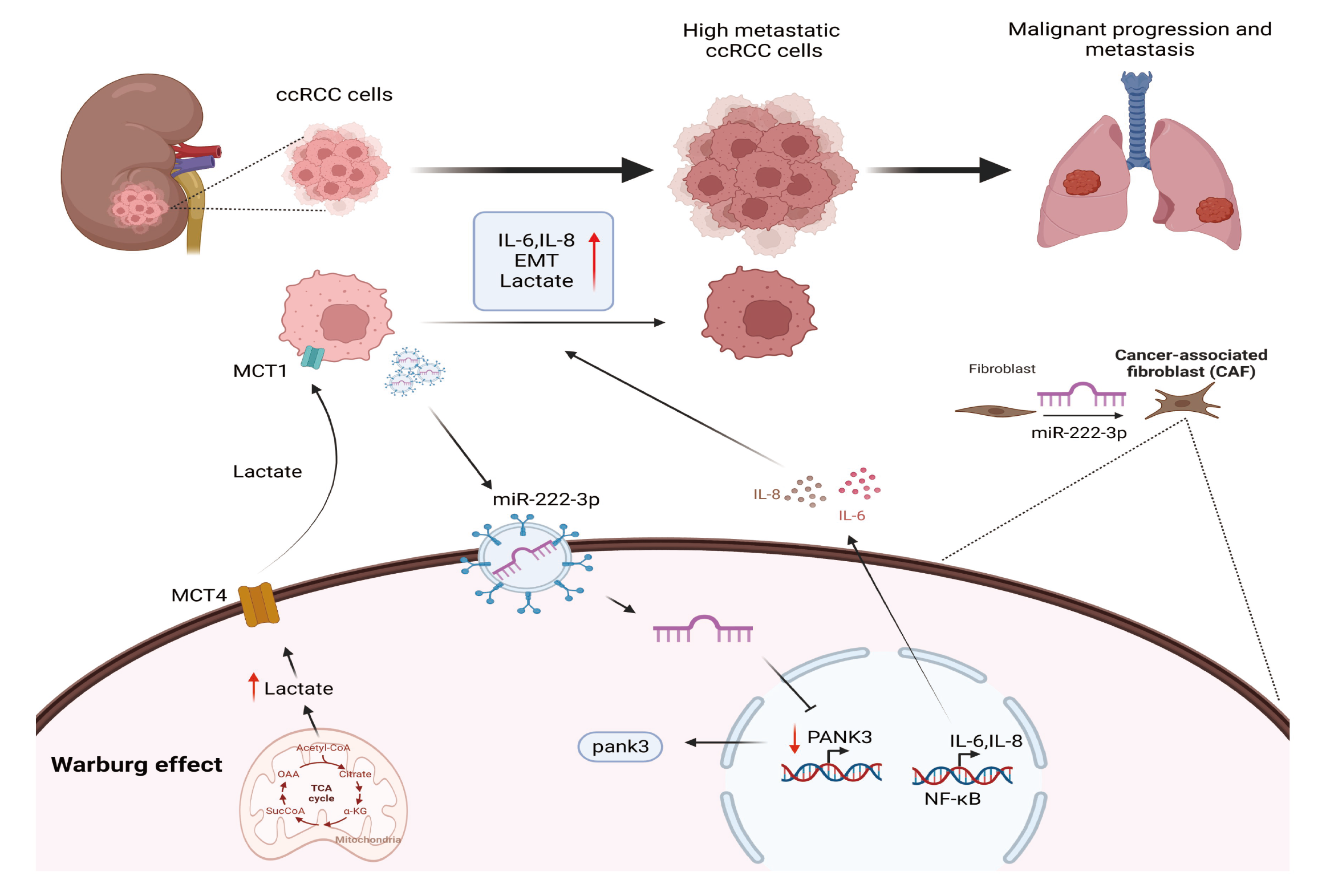
Proposed schematic diagram of tumor exosomal miR-222-3p-mediating fibroblasts activation to promote renal cancer progression.

In sum, our study demonstrates that exosomes derived from tumors contain miR-222-3p which activates the NF-κB signaling pathway leading to metabolic reprogramming. This is achieved by down-regulating PANK3, which transforms NFs into CAFs. The activated CAFs promote EMT and tumorigenicity in renal cancer cells by releasing pro-inflammatory factors and lactic acid. The findings suggest that exosomal miR-222-3p plays a significant role in intercellular communication, promoting the formation of an inflammatory microenvironment, and contributing to the progression of ccRCC.

## 4. Discussion

The tumor microenvironment is a complex system that relies on intercellular communication, and changes in this system are closely linked to tumor progression and metastasis^[29-30]^. Recent studies have shown that exosomes play a crucial role in intercellular communication^[19-20]^. Therefore, it is essential to investigate the interaction between exosome-mediated tumor cells and mesenchymal cells. In our study, we analyzed the differential exosomal-miRNA profiles of renal cell carcinoma cell lines with high and low metastatic backgrounds. Exosomes transfer miR-222-3p from tumor cells to fibroblasts, resulting in activation of the NF-kB signaling pathway in fibroblasts through down-regulation of the target gene PANK3 and metabolic reprogramming, which in turn activates CAFs. CAFs promote tumor progression by secreting inflammatory factors and lactic acid. This crosstalk between tumor cells and fibroblasts reveals the molecular mechanism of renal cancer progression. High expression of miR-222-3p in serum exosomes and ccRCC tissues is closely associated with renal cancer progression, highlighting the importance of effective prevention and treatment strategies.

Previous studies have reported abnormal expression of miR-222-3p in various tumors, with different effects. For instance, miR-222-3p can promote the proliferation, migration, and invasion of PTC cells by targeting SLC4A4^[31]^. Moreover, HBV infection can increase the expression of miR-222-3p, leading to the progression of HCC through miR-222-3p-mediated THBS1 downregulation^[32]^. Our findings reveal that tumor-derived exosomal miR-222-3p can activate fibroblasts in the microenvironment into CAFs. However, the role of miR-222-3p in the progression of renal cell carcinoma requires further investigation.

Coenzyme A is a vital cofactor in more than 70 enzyme reaction pathways in the body, providing 90% of the energy required for life, and also plays a crucial role in regulating glucose metabolism^[33-34]^. The study results demonstrate that the down-regulation of PANK3, mediated by miR222-3p, triggers the activation of fibroblasts possibly due to the activation of the NF-kB signaling pathway and metabolic reprogramming in fibroblasts. The down-regulation of coenzyme A promotes the aerobic glycolysis of NFs, leading to fibroblast activation through Warburg metabolism. The role of metabolic reprogramming in fibroblast activation requires further investigation; however, therapeutic strategies targeting metabolic reprogramming may provide a new way for cancer treatment.

CAFs have distinct differences in both structure and function when compared to fibroblasts^[10]^. It has been observed that the conditioned medium of CAFs can enhance the migration ability and EMT of renal cancer cells in vitro. Additionally, the conditioned medium and exosomes of CAFs have been found to promote the growth of renal cancer orthotopic transplantation tumor model and the formation of lung metastasis in nude mice in vivo, which is particularly noteworthy.

## 5. Conclusion

In sum, tumor-derived exosomal miR-222-3p is capable of transforming fibroblasts into CAFs by downregulating PANK3 expression in fibroblasts. This transformation leads to an increase in the secretion of inflammatory factors and lactic acid, which subsequently promotes tumor migration, proliferation, EMT, tumorigenicity, and lung metastasis. Furthermore, we have found a positive correlation between the high expression of miR-222-3p in serum exosomes and ccRCC tissues with the progression of renal cancer. Our findings shed light on a new molecular mechanism that explains how crosstalk between tumor cells and fibroblasts promotes the progression of ccRCC, and this can serve as a basis for developing effective prevention and treatment strategies for renal cancer.

## Authors’ Contributions

Yang Yang, Jie Zhu, Dong-lai Shen, Ce Han, Chen-feng Wang and Bo Cui contributed to performing the experiments, statistical analysis, article writing and editing. Wen-mei Fan, Xiu-bin Li, Yu Gao and Xu Zhang contributed to project design, article writing and editing.

## Compliance with ethical standards

### Ethical approval

Our study was approved by the local ethics committee and performed in accordance with the ethical standards of the institutional research committee.

### Informed consent

Informed consent was obtained from all individual participants included in the study.

